# Reciprocal zebrafish-medaka hybrids reveal maternal control of zygotic genome activation timing

**DOI:** 10.1101/2021.11.03.467109

**Authors:** Krista R. Briedis-Gert, Gunnar Schulze, Maria Novatchkova, Karin Panser, Luis Enrique Cabrera Quio, Anja Koller, Yixuan Guo, Bradley R. Cairns, Eivind Valen, Andrea Pauli

## Abstract

The sperm and egg contribute unequally to the newly formed zygote. While the sperm provides mainly paternal DNA, the egg provides both maternal DNA and the bulk of the future embryonic cytoplasm. Most embryonic processes like the onset of zygotic transcription are thought to depend on maternal cytoplasmic components, but this has not been tested rigorously. Here we report the establishment of a reciprocal zebrafish-medaka hybrid system which enables unequivocal distinction between maternal and paternal gene products. By combining expression of zebrafish Bouncer on the medaka egg with artificial egg activation, we demonstrate the *in vitro* generation of paternal zebrafish/maternal medaka (reripes) hybrid embryos. These hybrids complement the previously reported paternal medaka/maternal zebrafish (latio) hybrid embryos^1^, providing a versatile tool to dissect parental control mechanisms during early development. With this system, we investigated maternal vs. paternal control of zygotic genome activation (ZGA) timing. RNA-seq and ATAC-seq analyses of the purebred fish species and hybrids revealed that the onset of ZGA is primarily governed by the egg. Combining these datasets with proteome-wide analysis of early medaka and zebrafish embryogenesis highlights new potential regulators of ZGA, including Znf281b. Overall, our study establishes the reciprocal zebrafish-medaka hybrid system as a versatile tool to study parent-of-origin effects in vertebrate embryos.

## Introduction

Our limited ability to distinguish maternal and paternal components during development has hindered investigation into many fundamental mechanisms that govern embryogenesis. Researchers have long strived for an ideal system with which to accomplish this on the gene, transcript, and protein levels. Beyond studies at the single-gene level, a traditional approach has been to cross parents carrying different single nucleotide polymorphisms (SNPs)^2–4^. However, this methodology is restricted to the subset of loci containing SNPs. Thus, a hybrid system between two distantly related species would provide the unique advantage of combining highly dissimilar maternal and paternal genomes which could be unequivocally distinguished in their offspring.

Zebrafish and medaka are evolutionarily distant species separated by ∼110-200 MYR^5,6^. Both are well-established genetic models that can be housed in similar environmental conditions and undergo relatively fast early development. They are externally fertilizing and produce gametes and embryos that are amenable for live-cell imaging^5^. Importantly, zebrafish and medaka do not interbreed, and medaka sperm cannot fertilize zebrafish eggs *in vitro*^1^. Their genomes differ in sequence, size, and chromosome number. The medaka genome is 0.7 Gb, around half the size of that of zebrafish (1.5 Gb), and comprises 24 chromosomes including sex chromosomes, whereas the zebrafish genome constitutes 25 autosomes^7,8^. Despite these differences, both genomes contain a similar number of genes (∼25,000)^7,8^. Because even conserved genes differ in nucleotide sequence, most sequencing-derived reads from medaka-zebrafish hybrids should be unambiguously mappable to either genome. Thus, reciprocal hybrids from zebrafish and medaka would comprise a unique tool with which to address open questions of paternal and maternal control, conserved developmental mechanisms, as well as hybrid incompatibilities.

We previously reported that expressing the medaka homolog of Bouncer (Bncr), a factor essential for fish fertilization, on zebrafish eggs was sufficient to allow fertilization of these eggs by medaka sperm^1^. Following traditional hybrid nomenclature (paternal species x maternal species), we fused the species names of *Danio rerio* and *Oryzias latipes* to name the resulting hybrid embryos “latio” (male *O. **lat**ipes* x female *D. rer**io***). While latio hybrids alone are already a useful tool, the reverse hybrid, “reripes” (male *D. **rer**io* x female *O. lat**ipes***), has never been generated. The reripes hybrid would serve as the ideal complement to the latio hybrid as it would have an identical genome yet differ in the parent of origin for egg- vs. sperm-derived components. In this study, we report an efficient protocol to produce reripes hybrids and use the medaka-zebrafish reciprocal hybrid system to demonstrate maternal control of the timing of zygotic genome activation (ZGA).

## Results

We investigated whether reripes hybrid embryos could be produced in *in vitro* fertilization experiments with zebrafish sperm and medaka eggs overexpressing the zebrafish Bncr homology using a strategy similar to the one employed for generating latio hybrids^1^. While both fish are teleosts, medaka and zebrafish differ in their egg-laying behavior and produce telolecithal eggs with important physiological differences. Firstly, while unfertilized zebrafish eggs can be readily obtained from females *in vitro*^9^, efficient collection of unfertilized medaka eggs requires mating of medaka females with sterile males (see Methods)^10^. Secondly, in both organisms, the sperm contacts the egg only through a small opening in the protective envelope (chorion) around the egg. This opening, the micropyle, is a narrow canal through which the sperm must swim to reach the egg membrane. Because medaka and zebrafish sperm are similar in size^11–13^, the medaka micropyle should be able to accommodate zebrafish sperm. However, whether zebrafish sperm can indeed locate and pass through the medaka micropyle has not been tested. Thirdly, zebrafish sperm and eggs are activated upon contact with water, a hypotonic solution that induces sperm motility and renders the egg competent for fertilization^9,14^. In contrast, medaka eggs activate upon contact and/or fusion with sperm, while the sperm also acquires motility through contact with water^15^.

We first investigated whether zebrafish sperm can both locate and pass through the medaka micropyle. Live imaging of *in vitro* fertilization using MitoTracker-labeled zebrafish sperm and wild-type (medaka Bncr-expressing) as well as transgenic zebrafish Bncr-expressing medaka eggs revealed that zebrafish sperm are indeed able to find and enter the medaka micropyle independently of zebrafish Bncr expression (**Fig. 1A**). However, no zebrafish sperm were observed to enter the egg after reaching the micropylar pit, even after several minutes and irrespective of the expressed Bncr protein (**Fig. 1A**). In concordance with these observations, no fertilized eggs were obtained from *in vitro* fertilization (IVF) experiments with zebrafish sperm incubated with wild-type or zebrafish Bncr-expressing medaka eggs (**Fig. 1B**). In contrast, conspecific IVF of medaka sperm with wild-type medaka eggs resulted as expected in sperm-egg fusion after ∼2 minutes (**Fig. 1A**) and high fertilization rates (**Fig. 1B**). Thus, while expression of medaka Bncr on the zebrafish egg enables entry of medaka sperm resulting in latio hybrid embryos^1^, expression of zebrafish Bncr on medaka eggs is not sufficient for zebrafish sperm to enter medaka eggs.

**Figure 1.**
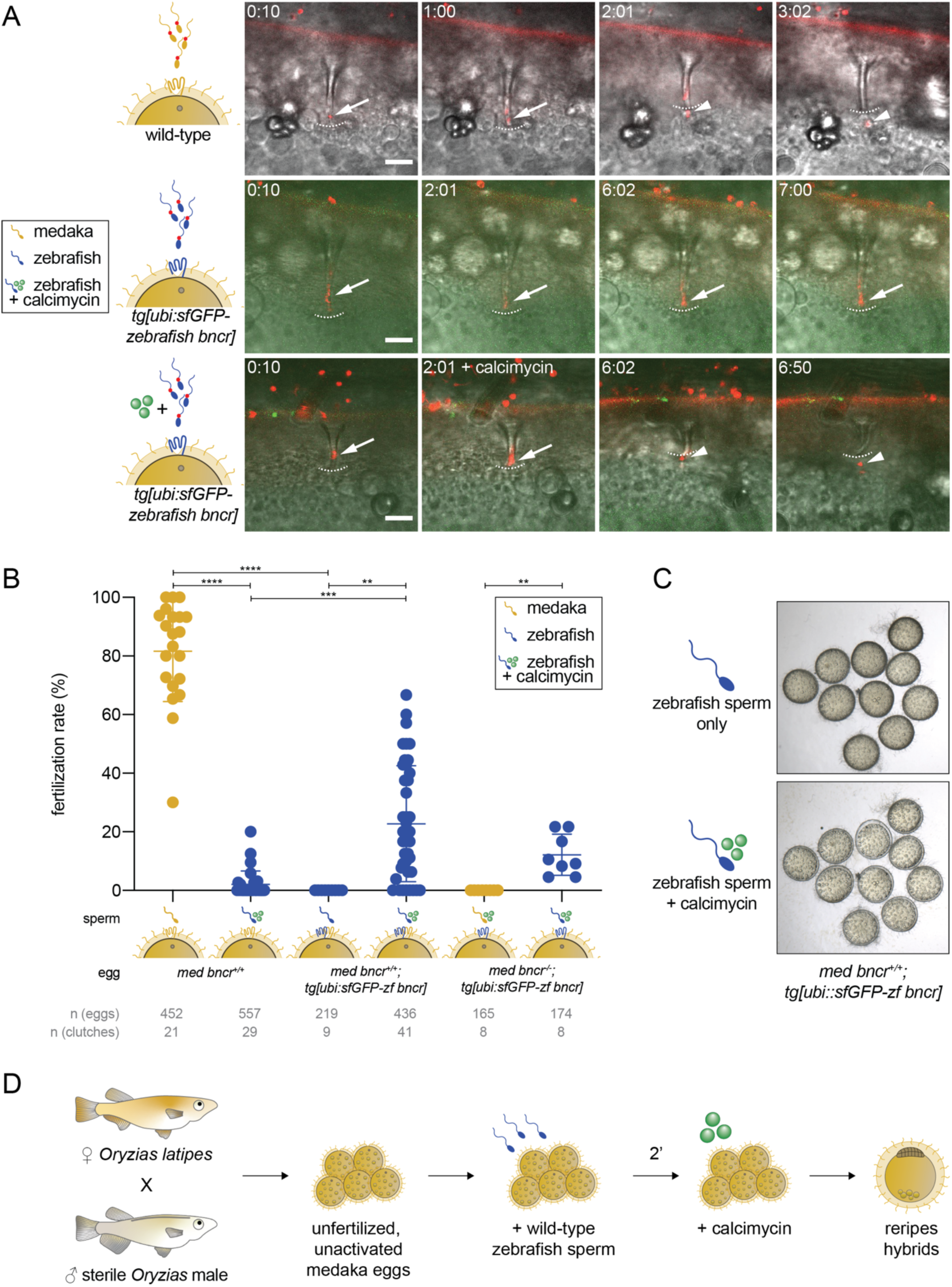
Zebrafish sperm can fertilize zebrafish Bncr-expressing medaka eggs upon artificial activation by calcimycin. **(A)** Time-lapse confocal images of MitoTracker-labeled sperm (red) incubated with medaka eggs. (Top) Medaka sperm efficiently fertilize wild-type medaka eggs and can be observed inside the egg ∼2 minutes post-sperm addition. (Middle) Zebrafish sperm find and enter the micropyle of medaka eggs expressing zebrafish Bncr (green), but do not fuse with the egg. (Bottom) Addition of calcimycin ∼2 minutes post-sperm addition induces egg activation and allows zebrafish sperm to fuse with zebrafish Bncr-expressing medaka eggs. White arrows indicate sperm inside the micropyle. Dashed white lines demarcate the base of the micropyle. White arrowheads indicate sperm inside the egg after it has fused with the egg membrane. Scale bar = 20 μm. **(B)** *In vitro* fertilization with medaka sperm and wild-type medaka eggs results in high fertilization rates (>60%), whereas medaka sperm cannot fertilize medaka *bncr*^-/-^ eggs expressing zebrafish Bncr. Zebrafish sperm are unable to fertilize zebrafish Bncr-expressing medaka eggs, but addition of calcimycin enables them to fertilize both wild-type and zebrafish Bncr-expressing medaka eggs in a wild-type or medaka *bncr*^-/-^ background. (IVF with *bncr*^+/+^ eggs, Kruskal-Wallis test with Dunn’s multiple comparisons test: ****adj. P = 1.08e-13 (medaka vs. zebrafish sperm + calcimycin on wild-type eggs), ****adj. P = 4.61e-9 (medaka sperm on wild-type eggs vs. zebrafish sperm on zebrafish Bncr-expressing eggs), ***adj. P = 1.36e-4, **adj. P = 0.005; IVF with *bncr*^-/-^ eggs, two-tailed Wilcoxon matched-pairs signed rank test: **P = 0.008). **(C)** Medaka eggs, even when expressing zebrafish Bncr, remain unactivated (top) in the presence of zebrafish sperm, as seen by the darker-colored cytoplasm and chorion tightly apposed to the egg membrane. Calcimycin addition induces artificial activation of medaka eggs (bottom), evident by the clear cytoplasm and gradual lifting of the chorion. **(D)** Workflow schematic depicting generation of reripes hybrids.

We observed that medaka eggs exposed to zebrafish sperm not only remained unfertilized, but also remained unactivated (**Fig. 1C**). We therefore hypothesized that zebrafish sperm are unable to activate medaka eggs and therefore fail to fertilize them even when they express zebrafish Bncr. Medaka eggs in calcium-containing media can be activated artificially using the calcium ionophore A23187 (calcimycin)^16,17^, a divalent cation ionophore that allows passage of calcium across membranes^18^. Remarkably, when zebrafish Bncr-expressing medaka eggs were incubated with zebrafish sperm followed by the addition of calcimycin, the eggs were both activated and fertilized (**Fig. 1A-C**). While fertilization success did not strictly depend on expression of zebrafish Bncr, fertilization rates were significantly higher with eggs expressing zebrafish vs. medaka Bncr, in line with the previously observed species specificity between medaka and zebrafish sperm and their species-matched Bncr proteins^1,19^ (**Fig. 1B**). Thus, reripes hybrids can be efficiently generated from zebrafish Bncr-expressing medaka eggs and zebrafish sperm upon artificial activation by calcimycin treatment (**Fig. 1D**).

The successful generation of reripes hybrids allowed us to compare development and gene expression in reripes and latio hybrids, which have identical genome content (haploid for both zebrafish and medaka genomes) yet differ in the origin of their maternal and paternal genomes and maternally provided cytoplasm (**Fig. 2A-B**). Overall, early development (up to 8 hours post-fertilization (hpf) of reripes hybrids resembles purebred medaka embryos, as was observed for latio hybrids and purebred zebrafish embryos (**Fig. 2A-B**). However, most reripes embryos arrest and begin to decompose before 16 hpf (**Fig. 2B**), coinciding with the time of gastrulation^20^. Thus, reripes hybrids survive for a shorter period of time than latio hybrids, which arrest only during somitogenesis and start to decompose ∼24 hpf^1^ (**Fig. 2A**).

**Figure 2.**
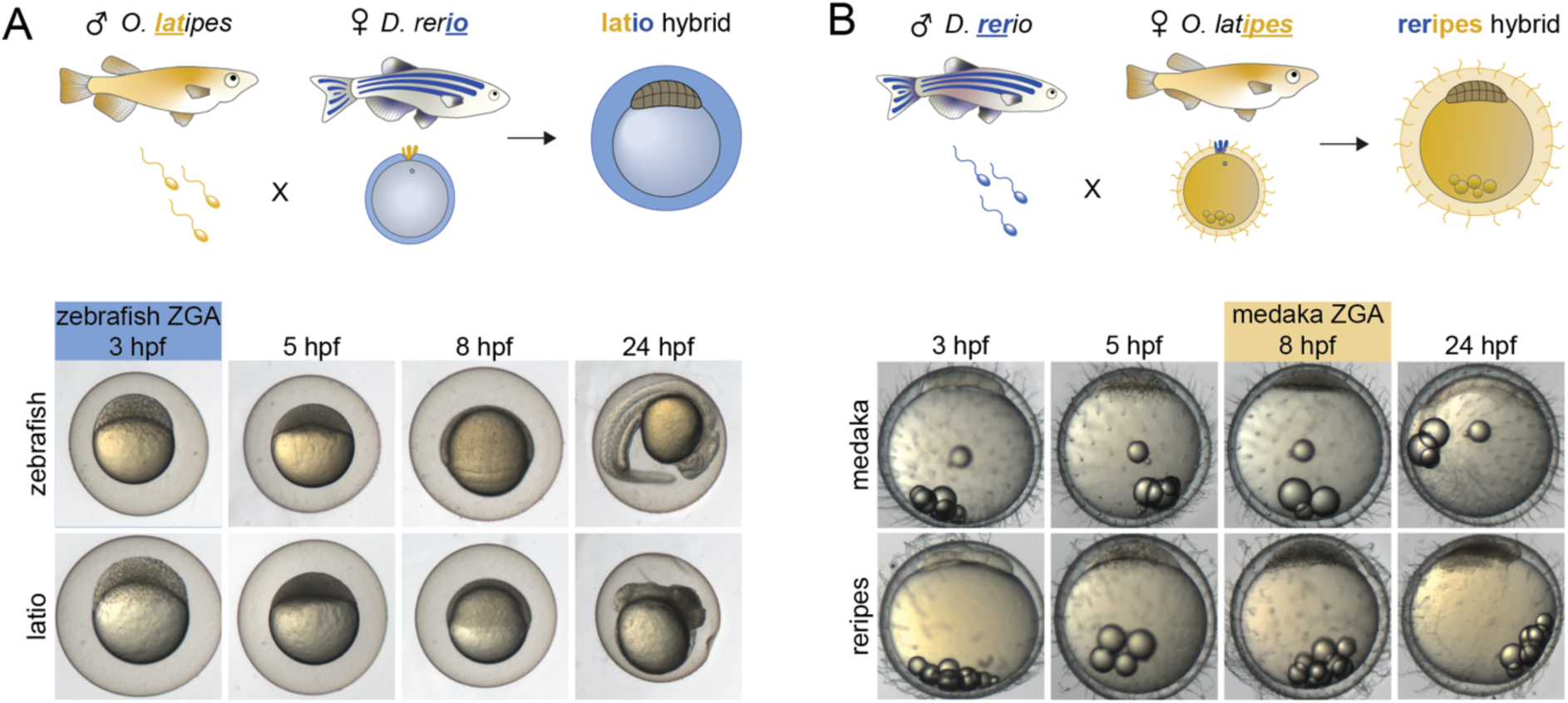
Development of latio and reripes hybrids resembles that of the maternal fish species. **(A)** Schematic depicting the naming convention for latio and reripes hybrids, whose names are derived from fusing the paternal and maternal species epithets, respectively. **(B)** Bright-field image time course of zebrafish, latio, medaka, and reripes embryos. Overall, latio hybrids resemble zebrafish embryos, while reripes hybrids resemble medaka embryos. However, in both cases, a delay evident by morphology of the hybrid embryo is apparent by 8 hpf. While latio hybrids undergo gastrulation, both hybrids have arrested development and are necrotic by 24 hpf.

Despite hybrid inviability in both crosses, latio and reripes hybrids develop past the time of ZGA in both parental species. The major wave of ZGA occurs at ∼3 hpf in zebrafish^21,22^, while medaka ZGA takes place later at ∼6-8 hpf^23,24^ (**Fig. 2A-B**). This intrinsic difference between medaka and zebrafish development presented us with a unique opportunity to investigate maternal vs. paternal control of ZGA timing. In the medaka-zebrafish hybrid system, the same genome exists in either an early-activating (zebrafish) or a late-activating cytoplasm (medaka). To determine whether the foreign paternal genome is expressed in both hybrids and to compare gene expression dynamics in purebred vs. hybrid embryos, we performed ribosomal RNA-depleted RNA-sequencing of zebrafish, medaka, latio, and reripes samples collected at 3, 5, 8, and 24 hpf (no 24-hpf sample was collected for reripes given their arrest at ∼16 hpf). Consistent with maternal contribution of mRNAs, reads from the earliest time point mapped to the corresponding maternal genome in both hybrids (**Fig. 3A**). However, reads from later time points mapped uniquely to regions in zebrafish and medaka genomes for both hybrid samples (**Fig. 3A**), demonstrating that the zebrafish genome is expressed in the medaka egg (reripes hybrid) and the medaka genome is expressed in the zebrafish egg (latio hybrid). Although only a small fraction of the total number of reads was derived from the paternal genome during the first hours after ZGA onset in both hybrids (**Fig. S1A**), thousands of paternal genes became expressed at the latest time point analyzed for each hybrid, as evidenced by the ∼15,000 single-species protein-coding genes expressed in purebreds across all time points vs. ∼30,000 zebrafish and medaka protein-coding genes expressed in late hybrid samples (**Fig. 3A, Table S1**). Therefore, we conclude that genome activation in zebrafish-medaka hybrids occurs genome-wide and that both zebrafish and medaka cytoplasmic components can drive successful activation of the other species’ genes.

**Figure 3.**
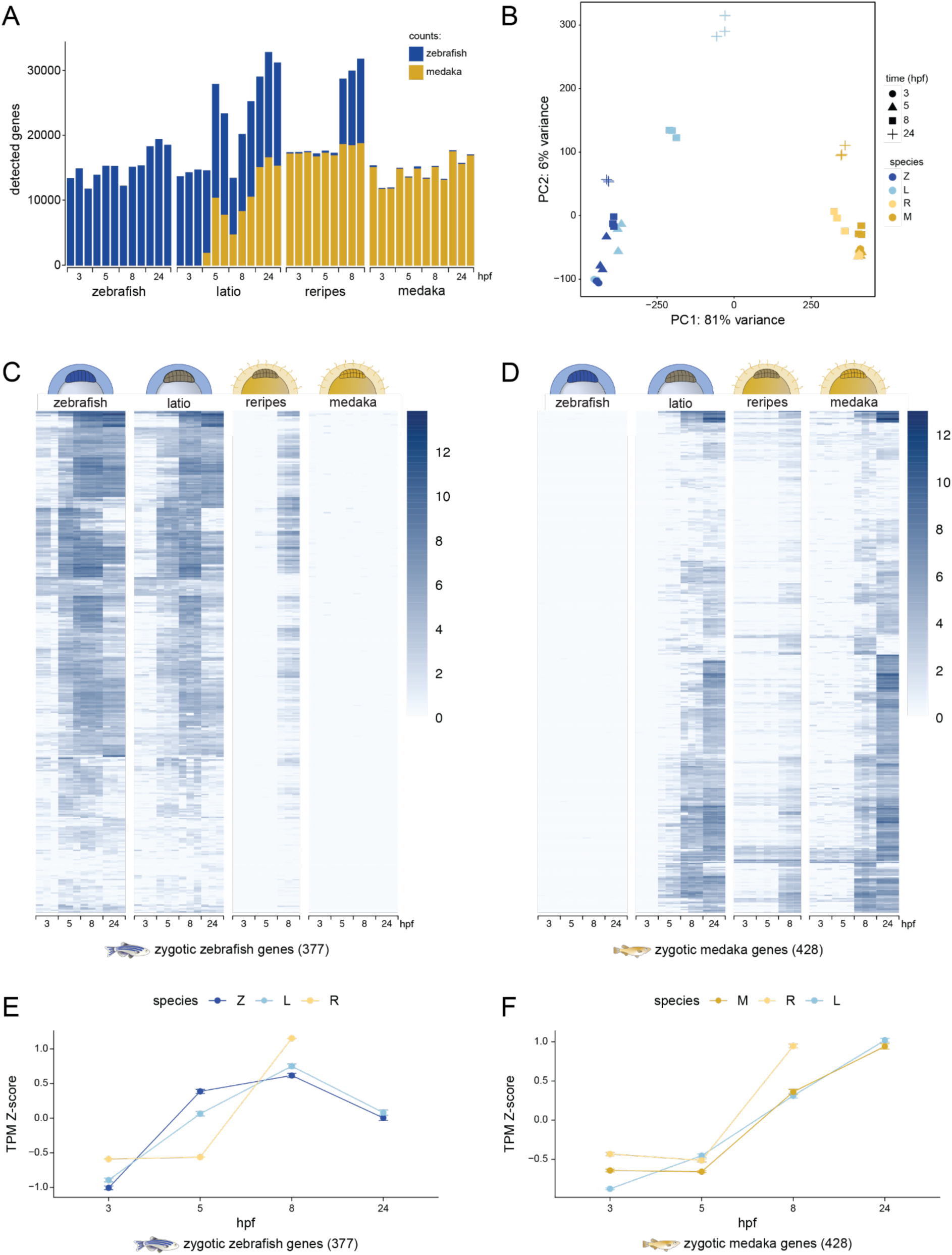
The foreign paternal genome is expressed and activated according to the egg’s timing in both hybrids. **(A)** Number of protein-coding genes detected (read counts >3) in each sample with genome of origin indicated (zebrafish in dark blue, medaka in dark yellow). Upon expression of the foreign paternal genome, the number of detected genes nearly doubles in latio and reripes as a result of both the medaka and zebrafish genomes’ being expressed. (**B**) Principal Component Analysis (PCA) of the top 1000 differentially expressed genes per sample. **(C)** Heat map of gene expression dynamics in zebrafish, latio, reripes, and medaka embryos for the subset of 377 protein-coding genes that are purely zygotically expressed in purebred zebrafish. Expression values are plotted as log2(TPM). **(D)** Heat map of gene expression dynamics in zebrafish, latio, reripes and medaka embryos for the subset of 428 protein-coding genes that are zygotically expressed in purebred medaka. Expression values are plotted as log2(TPM). (**E, F**) TPM Z-score plots of zygotic zebrafish (**E**) and zygotic medaka (**F**) genes. Zebrafish zygotic genes are delayed in reripes hybrids, while medaka zygotic genes are prematurely activated in latio hybrids.

Because medaka and zebrafish naturally differ in the time of ZGA onset and both hybrids actively express both genomes, we analyzed the timing of ZGA in latio and reripes hybrids. Principle Component Analysis (PCA) of the top 1000 differentially expressed genes amongst all samples revealed that hybrid embryos clustered at early time-points with their maternal species, but started to diverge after the onset of ZGA of the respective maternal species (**Fig. 3B**). To examine gene expression at ZGA in more detail, we limited the analysis to purely zygotically expressed genes (excluding maternal-zygotic genes), which we defined based on embryonic gene expression time courses of zebrafish (by SLAM-seq^22^) and medaka (by RNA-seq^25^) (see Methods). Comparison of the onset of ZGA in hybrid embryos vs. parental species revealed that maternally and paternally derived genes became activated at similar times in a given hybrid embryo and followed the maternal timing (**Fig. 3C-F**). As such, robust onset of paternal zebrafish gene expression was detected in the reripes hybrid at 8 hpf, coinciding with the onset of expression of zygotically expressed medaka genes, yet delayed compared to the normal timing of zebrafish ZGA at ∼3 hpf (**Fig. 3C, E**). Along the same lines, onset of paternal medaka gene expression was already detected at 5 hpf in the latio hybrid, which is premature in comparison to ZGA timing in medaka at ∼8 hpf (**Fig. 3D, F**).

To compare the expression dynamics of the zebrafish and medaka alleles for a given gene directly, we extended our analysis to the subset of orthologous gene pairs between zebrafish and medaka. In hybrid embryos, both zebrafish and medaka alleles of orthologous genes were generally activated simultaneously (**Fig. S1B-C**, **Table S2 and S3**). (Note that a subset of orthologous genes that are zygotically expressed in zebrafish are maternally provided in medaka, which precluded their analysis. For example, in latio hybrids, onset of expression of the paternal medaka ortholog at ∼5 hpf temporally matched that of the maternal zebrafish ortholog at ∼5 hpf, whereas the same medaka allele would be expressed only at ∼8 hpf in reripes hybrid and purebred medaka embryos (**Fig. S1B-C**). Thus, despite having identical genome content, reripes and latio embryos activate their genomes at different times, providing direct evidence that the timing of ZGA is largely influenced by factors in the egg and is not intrinsic to paternal chromatin.

Chromatin accessibility is directly linked to gene expression. Based on the observation that ZGA timing correlates with the intrinsic timing of the maternal cytoplasm, we investigated whether chromatin accessibility could explain the shifted timing of paternal gene expression in the hybrids. We hypothesized that in the medaka cytoplasm (reripes hybrids), zebrafish chromatin maintains a closed state for a prolonged period of time until its expression at ∼8 hpf, leading to its delayed activation. Conversely, we hypothesized that in the zebrafish cytoplasm (latio hybrids), medaka chromatin becomes accessible earlier, enabling premature expression of zygotic genes at ∼5 hpf. Alternatively, the paternal zebrafish genome in the reripes hybrid may become open early relative to the medaka genome, but gene expression could be delayed due to insufficient levels of transcriptional activators in the medaka egg.

To investigate the timing of chromatin opening, we performed ATAC-seq (Assay for Transposase-Accessible Chromatin using sequencing) of purebred medaka and reripes hybrid embryos at 5 and 8 hpf and purebred zebrafish and latio hybrid embryos at 3, 5, and 8 hpf. Genes were classified into three sets based on their expression status in purebreds and hybrid embryos (**Fig. 3A, Table S1**): (1) zygotic genes are those that are zygotically expressed in purebred species excluding maternal-zygotic genes; (2) paternal zygotic genes are those that are zygotically expressed from the paternal genome in hybrid embryos and not included in set (1); this is the largest group of genes and includes maternal-zygotic as well as unexpressed genes in purebred embryos; (3) unexpressed genes are those that are not expressed in either purebred or hybrid condition during the time course analyzed here.

Overall, the zebrafish genome exhibits a higher degree of accessibility near transcription start sites (TSSs) (± 50 bps) of both zygotic gene sets compared to those in the medaka genome, independently of the parent of origin and even at time points when zygotic transcription occurs in both respective genomes (**Fig. 4A-B**). Paternal zygotic genes were found to increase in accessibility at the same time yet to a slightly larger extent compared to zygotic ones, and ∼65% were found to be maternal-zygotic genes based on SLAM-seq data^22^. This suggests that their expression during oogenesis may prime them for efficient opening during embryogenesis^23^. In line with genome accessibility timing correlating with the intrinsic timing of the maternal cytoplasm, accessibility of the zebrafish genome was greatly reduced in the medaka cytoplasm compared to its own cytoplasm (reripes at 5 hpf vs. zebrafish and latio at 5 hpf) (**Fig. 4B**). Even at 8 hpf when accessibility had increased in reripes hybrids, accessibility of the zebrafish genome was lower than in latio and zebrafish embryos at 3 and 5 hpf. Conversely, the medaka genome became accessible earlier in the zebrafish cytoplasm (latio at 3 hpf) compared to both medaka and reripes at 5 hpf (pre-ZGA) and to the same level as the maternal zebrafish genome (Fig. **4B**). Thus, onset of zygotic gene expression from the foreign (paternal) genome correlates with an increase in chromatin accessibility at TSSs of zygotically expressed genes (**Fig. 4C**).

**Figure 4.**
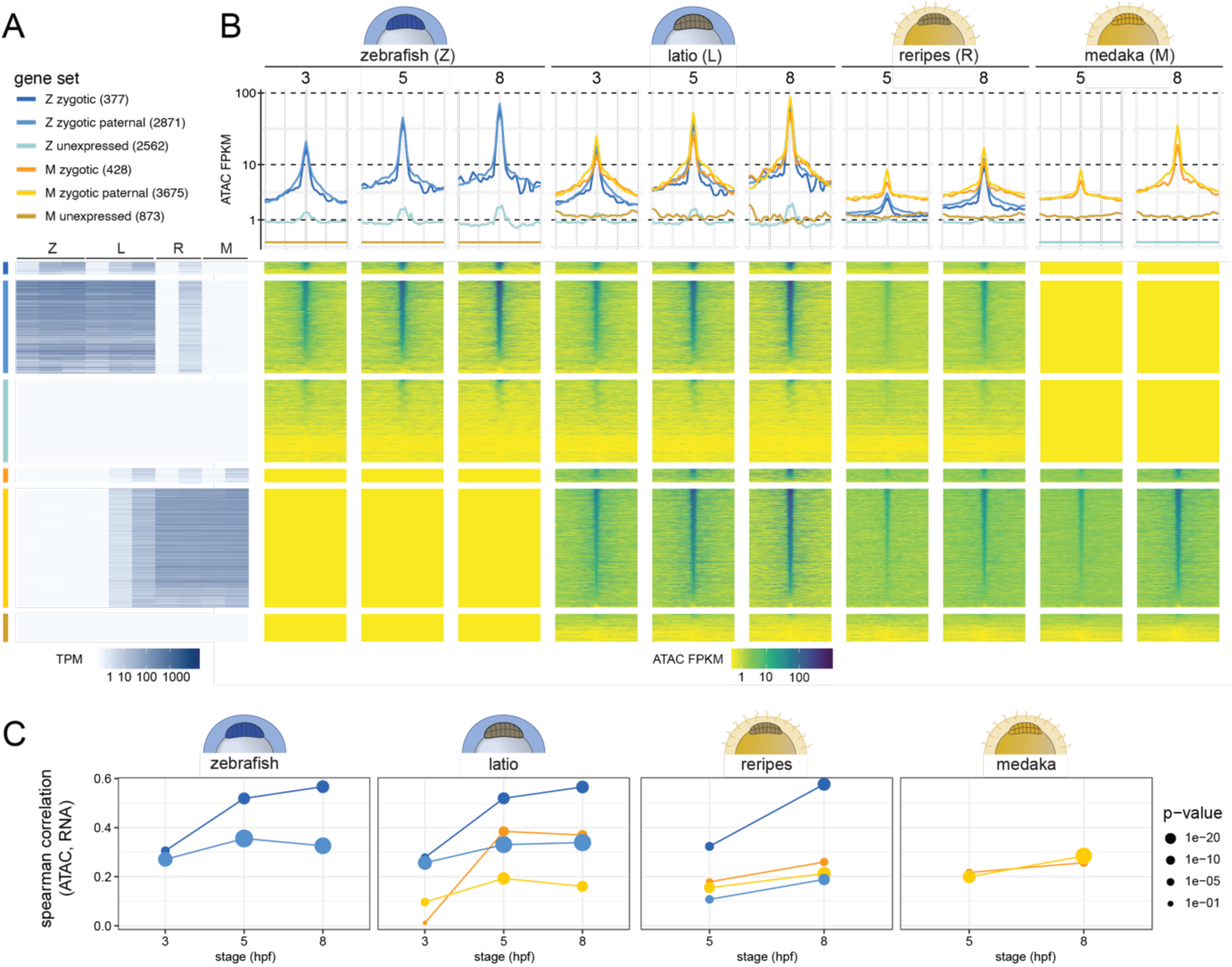
Accessibility of the paternal genome mirrors that of the maternal genome in hybrids. **(A)** Gene sets analyzed in ATAC-seq. Zygotic gene sets were defined by SLAM-seq and RNA-seq data (see Methods). Unexpressed genes are those without any RNA-seq read counts over the time course (up to 24 hpf). Heat map of the gene expression patterns (RNA-seq) corresponding to the genes plotted on the right. TPM, transcripts per million. Z = zebrafish, M = medaka, L = latio, R = reripes. **(B)** Top: Metaplot of mean ATAC-seq signal normalized per genome around TSSs (± 2-kb window) for each gene set. Bottom: Heat map of ATAC-seq signal normalized per genome for 100-bp bins around TSS (± 2-kb window) for each gene set. Genes are sorted by accessibility in the center bin (TSS ± 50 bp) at 5 hpf for zebrafish and 8 hpf for medaka genes. Gene categorizations for each heatmap are shown in (A). FPKM, fragments per kilobase per million mapped reads. **(C)** Spearman correlation plots for ATAC-seq and RNA expression values. For purebreds, values are only plotted for gene sets that are expressed (gene sets 1 and 2).

As both ZGA timing and chromatin accessibility were found to depend on the maternal cytoplasm, we sought to identify candidate timing factors in the medaka egg whose expression dynamics differed temporally and/or in magnitude compared to the zebrafish homolog. To this end, we analyzed sequences corresponding to regions significantly increasing in accessibility (log2 fold change >1, FDR <5%) in each species’ ZGA timeframe (from 3 to 5 hpf in zebrafish and 5 to 8 hpf in medaka) (**Fig. 5A-B**). Comparing sequences from these differentially accessible regions (DARs) with those from regions without changes in accessibility, we identified transcription factor motif clusters specifically enriched in DARs in zebrafish and medaka (**Fig. S2A**). As expected, and in line with prior studies^26–29^, individual motifs among the top ten most highly enriched in zebrafish DARs corresponded to Pou5f3 (Pou5f1/Pou2f1), Sox19b (Sox2), Nanog, and Znf281, as well as to the zinc finger transcription factor Snai3 (**Fig. 5C, Fig. S2A**). In medaka, motifs for Nanog, Pou5f3/Oct4, Sox2, and Znf281 were also enriched in DARs, but not as highly as those corresponding to members of the Sp1 C2H2-type zinc-finger protein family, Klf15 and Sp2, as well as Patz1, Zbtb14, Tcfl5, Elk3, Etv5, and the forkhead box proteins Foxj3 and Foxn3 (**Fig. 5C, Fig. S2A**).

**Figure 5.**
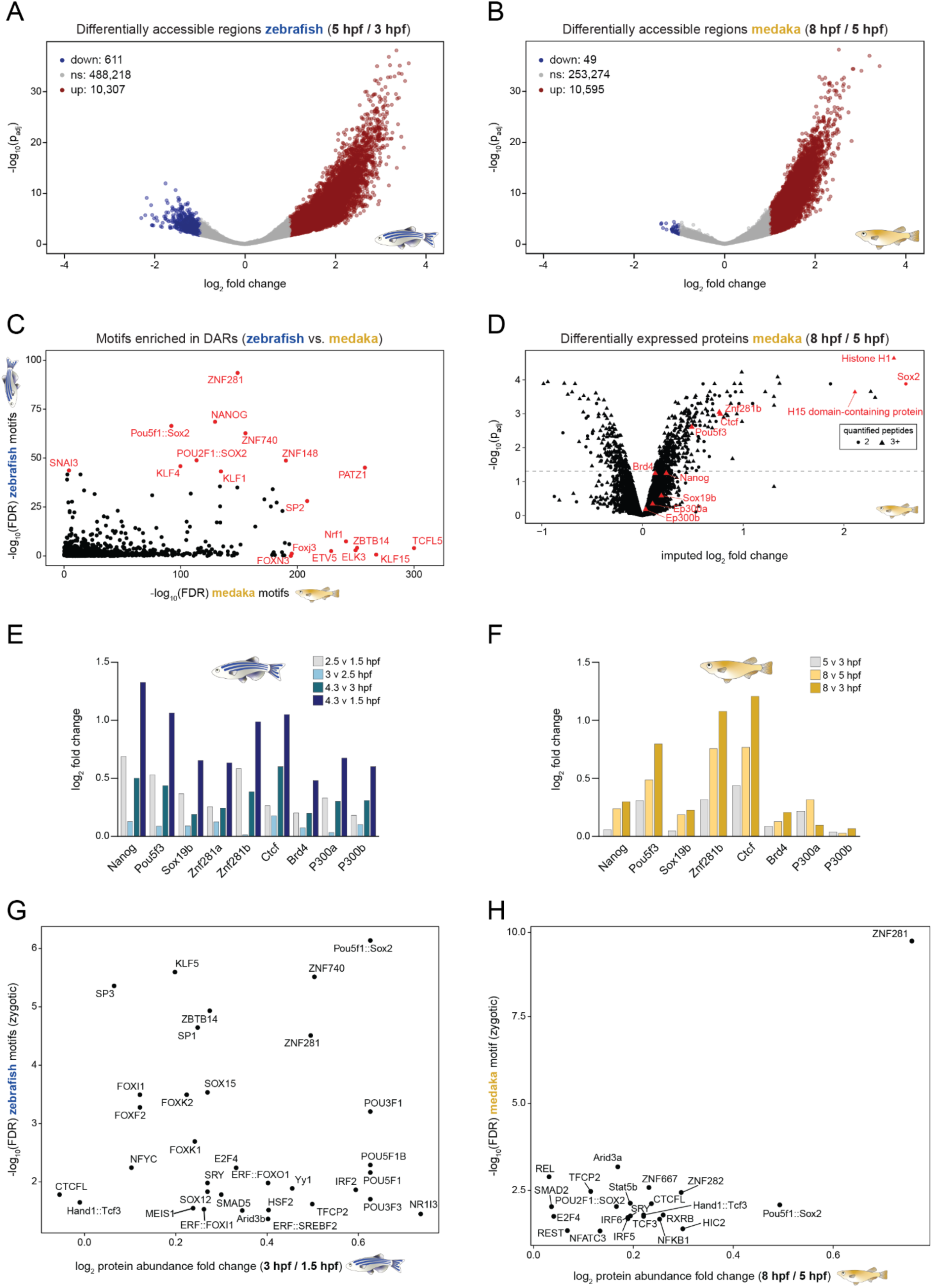
Identification of new potential regulators of ZGA. **(A-B)** Volcano plots for ATAC-seq peaks in differentially accessible regions (DARs) that are either decreasing (down, blue), unchanged (gray), or increasing (up, red) in accessibility in zebrafish from 3 to 5 hpf or in medaka from 5 to 8 hpf. **(C)** Peak regions significantly increasing in accessibility were used for motif enrichment analysis (see Methods). Transcription factor motifs enriched in zebrafish DARs vs. medaka DARs are plotted. The top 10 motifs in zebrafish and/or medaka DARs are highlighted. **(D)** Differentially upregulated and downregulated proteins detected in the TMT-MS dataset of medaka (8 hpf versus 5 hpf). Transcription factors and other proteins previously implicated in ZGA and/or found to be enriched based on both motif analysis and on the protein level are highlighted in red. **(E-F)** Log2 fold changes of candidate transcription factors and chromatin remodelers in zebrafish and medaka from TMT-MS datasets. Of the candidates shown, only Pou5f3/Oct4, Ctcf, and Znf281b increase comparably leading up to ZGA at 8 hpf in medaka, whereas Nanog, Pou5f3, Sox19b, and Znf218b show the highest increase pre-ZGA (2.5 v 1.5 hpf) in zebrafish. **(G-H)** Motif enrichment (-log10(FDR)) for differentially accessible promoter regions in zygotic zebrafish genes versus zebrafish TMT protein expression values of the corresponding transcription factor (G); the corresponding plot for medaka is shown in (H).

To quantify the protein dynamics of transcription factors corresponding to these motifs, we performed tandem mass tag mass spectrometry (TMT-MS) of wild-type medaka embryos at 3, 5, and 8 hpf (**Fig. 5D**) and compared both absolute protein levels (**Fig. S2B-C**) and fold changes over time to an existing zebrafish TMT-MS dataset^30^ (**Fig. 5E-F**). Of the transcriptional regulators observed to increase significantly between 5 and 8 hpf in medaka (**Fig. 5D, F**), we found Znf281b to be among those with the highest fold change. Ctcf and Pou5f3/Oct4 were also observed to increase significantly over time, with Nanog and Sox19b showing increasing but lower fold changes during the same timeframe (**Fig. 5D, F**). Comparing their dynamics in medaka vs. zebrafish, both Pou5f3/Oct4 and Znf281b increase prior to or during ZGA in both species, whereas Nanog and Sox19b show a sharp increase pre-ZGA in zebrafish embryos but not in medaka (**Fig. 5E-F**, **Fig. S2B-C**). We then filtered the enriched motifs for only those present in zygotic gene promoters in each species (**Fig. 4A**, “Z zygotic” and “M zygotic”; **Table S1**) and only those with corresponding TFs detected in TMT-MS (**Fig. 5G-H, Fig. S2D-E**). Pou5f1/Sox2, Znf740, and Znf281 motifs were among the most significantly enriched motifs in zebrafish zygotic gene promoters that also had the highest fold changes in corresponding TF protein expression from 1.5 to 3 hpf in TMT-MS (**Fig. 5G**). In medaka zygotic gene promoters, the Znf281 motif was the single topmost enriched and its corresponding TF Znf281b was the most highly upregulated TF protein during ZGA (**Fig. 5H, Fig. S2E**). Importantly, the Znf281 motif was found in 68% of zygotic medaka genes compared to 24% of zebrafish zygotic genes. Thus, based on its robust motif enrichment in accessible zygotic gene promoters and strong upregulation on the protein level leading up to ZGA, Znf281b may play an important role in the temporal regulation of ZGA in medaka. Similarly, Znf740 may also represent a so-far unappreciated factor in zebrafish ZGA given its motif enrichment in zygotic genes and large increase on the protein level pre-ZGA. Together, these results suggest the presence of divergent gene regulatory mechanisms in zebrafish and medaka that, while different, maintain the ability to activate the other species’ genome in the context of ZGA.

## Discussion

Complementary latio and reripes hybrids constitute a distinctive system for studying early development with the advantage of clear differentiation between maternal and paternal components in the embryo. The generation and initial characterization of these hybrids presented here provide insights into both the processes of egg activation and fertilization as well as the control of ZGA timing. We envision this hybrid system as a tool with widespread applications, from studying inheritance of maternal piRNAs^31^ or epigenetic marks^32^, to further probing the intricacies of transcriptional activation during embryogenesis.

Fertilization of zebrafish Bncr-expressing medaka eggs by zebrafish sperm extends and strengthens our previous finding that Bncr is sufficient to mediate sperm-egg compatibility in zebrafish and medaka^1^. However, despite Bncr’s conserved role, zebrafish and medaka have diverged in their modes of egg activation. While zebrafish sperm binding via Bncr is not sufficient to trigger medaka egg activation, this can be induced artificially using the calcium ionophore calcimycin. We speculate that PLCZ1, a sperm-provided factor that induces mammalian egg activation via the release of calcium from internal stores in the endoplasmic reticulum^33–35^, may be required in medaka, but not zebrafish. Consistent with this idea, *plcz1* is expressed in medaka testis^36^ but transcripts of the closest zebrafish homolog based on protein sequence (*plcd4a*) are absent from zebrafish testis in published RNA-seq data^1,37^.

In addition to the different requirements for egg activation in zebrafish vs. medaka, latio and reripes hybrids revealed differences in length of survival and developmental activation of the zygotic genome dependent on the orientation of the cross. Latio hybrids survive for a longer period of time than reripes hybrids (24 vs. <16 hours), highlighting asymmetry in the ability of each species’ cytoplasm to cope with a haploid foreign genome. Such a phenomenon has been previously observed for paternal *Xenopus tropicalis* x maternal *X. laevis* hybrids which are viable, while the reverse cross fails to reach gastrulation^38,39^.

The juxtaposition of two differently timed genomes within the same cytoplasm in both parental orientations allowed us to dissect the roles of paternal and maternal contributions in determining ZGA timing. Our results reveal that the paternal genome does not retain an intrinsic species timing for gene expression when placed into a foreign cytoplasm, but instead adopts the timing imposed by the maternal cytoplasm (**Fig. 3**). This is further evident by the fact that even though latio and reripes hybrids are genetically identical, they undergo ZGA at different times, providing direct evidence for cytoplasmic rather than genomic control of ZGA timing during embryogenesis. In the same way, we observed that the paternal genome adopted the timing of increase in chromatin accessibility shown by the maternal genome in both hybrids, suggesting that factors controlling this accessibility likely contribute to ZGA timing (**Fig. 4, 5**).

The innate differential ZGA timing in medaka and zebrafish begs the question of what cytoplasmic factors drive this difference. Several models have been proposed to explain what regulates ZGA timing, including lengthening of the cell cycle to allow sufficient time for transcription, titration of maternal repressors (such as histones) as the nucleocytoplasmic (N/C) ratio increases, or an increase in nuclear concentration of specific transcriptional activators over time^40–43^. Our observations in the hybrids argue against a major role for N/C ratio and cell cycle length as the main regulators of ZGA timing. In the reripes hybrid, the N/C ratio is increased relative to a medaka embryo, and cell cycle length is increased relative to a zebrafish embryo, which would be predicted to shift ZGA earlier in both cases. Nevertheless, ZGA occurs normally for medaka genes, and zebrafish genes are activated much later (∼8 hpf) than in a zebrafish embryo (∼3 hpf) (**Fig. 3A, C-D**).

From the perspective of competition between histones as maternal repressors and transcription factors as maternal activators of transcription^44^, one could speculate that the early-activating zebrafish cytoplasm tips this balance in favor of transcription sooner than medaka, allowing earlier ZGA. In zebrafish, it has been shown that the transcription factors Pou5f3, Sox19b, and Nanog are required for activating expression of the earliest zygotic genes^26,27^ and that the concentration of free histones in the nucleus decreases with ZGA onset^44^, thus allowing transcription factor binding. Transcriptional competency of the earliest transcribed genes has been shown to be mediated by H3K27Ac writing and reading by P300 and Brd4, respectively, demonstrating the importance of chromatin accessibility in this context^45^. Analogous studies have not yet been done in medaka, yet we find that while Brd4, P300a, and P300b do increase in the medaka embryo prior to ZGA, their increase is not as pronounced as it is in the zebrafish embryo pre-ZGA (**Fig. 5E-F**). In medaka, the histone/transcription factor balance may be shifted to favor later onset of ZGA. We find that two histone proteins (histone H1 and H15 domain-containing protein) are significantly upregulated prior to ZGA onset, potentially prolonging the time needed to tip this balance in favor of zygotic transcription (**Fig. 5D**). It has been shown, however, that RNA polymerase II is phosphorylated in most cells of the medaka embryo at approximately 4 hpf (128 cells) and that cell divisions lose their synchrony prior to the mid-blastula transition (in contrast to zebrafish)^46^, suggesting that additional factors likely differentiate medaka developmental timing from that of zebrafish.

Previous studies of the role of Nanog and Oct4/Pou5f3 in medaka have already implied species-specific differences for their roles in early development between medaka and zebrafish, supporting the idea that medaka may rely on alternative factors to initiate ZGA^47,48^. In this study, we find that Znf281b motifs are among those most highly enriched in regions of increasing accessibility in the medaka genome during ZGA (**Fig. 5C, H**). Moreover, Znf281b protein levels increase significantly from 5 to 8 hpf in medaka (**Fig. 5D, F**), raising the possibility that this transcription factor may play a key and previously overlooked role in the induction of zygotic gene expression in medaka. While Znf281b motifs were not as highly enriched in open chromatin in zebrafish (**Fig. 5G**), we found that zebrafish Znf281b protein levels also increase pre-ZGA similarly to Nanog, Pou5f3, and Sox19b (**Fig. 5E**), which is consistent with what has recently been reported in zebrafish^29^. In that study, CRISPR-Cas13d mediated knockdown of *znf281b* mRNA in zebrafish resulted in a developmental delay during embryogenesis, suggesting an important function during early development by potentially regulating zygotic transcription with Nanog^29^. Future studies will be required to determine whether Znf281b is required for ZGA in medaka and how it functions mechanistically. Identifying which genes are regulated by this factor will elucidate the impact it may have on embryonic processes including and beyond ZGA. Given the large number of genes with Znf281b motifs in their promoter regions, our data raises the possibility that it could have a major influence on ZGA and subsequent development particularly in medaka. Intriguingly, studies in mouse embryonic stem cells have revealed an interconnected pluripotency network consisting of NANOG, OCT4, and ZFP281, the mouse ortholog of Znf281b, and provide evidence for direct interaction among these factors in regulating gene expression^49–52^. ZFP281 has further been shown to be required for transcription of NODAL signaling genes, acting in concert with OCT4 and P300 during mouse embryogenesis *in vivo*^53^.

Overall, this study demonstrates the generation of a novel reciprocal hybrid system between medaka and zebrafish, facilitating future studies that may uncover deeply conserved embryonic programs between these two evolutionarily distant species. In addition, this system will enable future work on unraveling the molecular basis for developmental allochrony, as medaka and zebrafish exhibit salient temporal differences in overall development including ZGA onset.

## Author Contributions

KRBG and AP conceived the study; KRBG designed, performed, and analyzed experiments; GS, MN and LECQ performed computational analyses of the RNA-seq data, and GS and MN analyzed ATAC-seq and proteomics data; KP and AK assisted with hybrid and ATAC-seq experiments, respectively; YG and BRC contributed to RNA-seq data generation; EV supervised and contributed to computational analyses; AP coordinated and supervised the project; KRBG and AP wrote the manuscript with input from all authors.

## Acknowledgments

We thank Katharina Lust (IMP Vienna, Austria) for providing a plasmid and valuable advice for working with medaka; Kiyoshi Naruse (Okazaki, Japan) for helpful advice, and the National Institute for Basic Biology (NIBB) (Okazaki, Japan) for providing *O. curvinotus*; the team of the BioOptics facility at the Vienna Biocenter, in particular Pawel Pasierbek for support with microscopy; the Next Generation Sequencing facility at the Vienna BioCenter Core Facilities (VBCF) for sequencing; the Proteomics Facility, in particular Michael Schutzbier, Richard Imre, and Elisabeth Roitinger for performing TMT-MS analysis of medaka samples; the animal facility personnel from the IMP for taking excellent care of zebrafish and medaka; and the entire Pauli lab, particularly Victoria Deneke and Jessica Stock, for fruitful discussions.

## Data availability statement

The original contributions presented in the study are included in the article/supplementary material; RNA-Seq data from this study were deposited at Gene Expression Omnibus under accession number GSE204802; ATAC-seq data from this study were deposited at Gene Expression Omnibus under accession number GSE314827; all inquiries should be directed to the corresponding author.

## Supplementary Tables & Figures

### Supplementary Figures

**Supplementary Fig S1.**
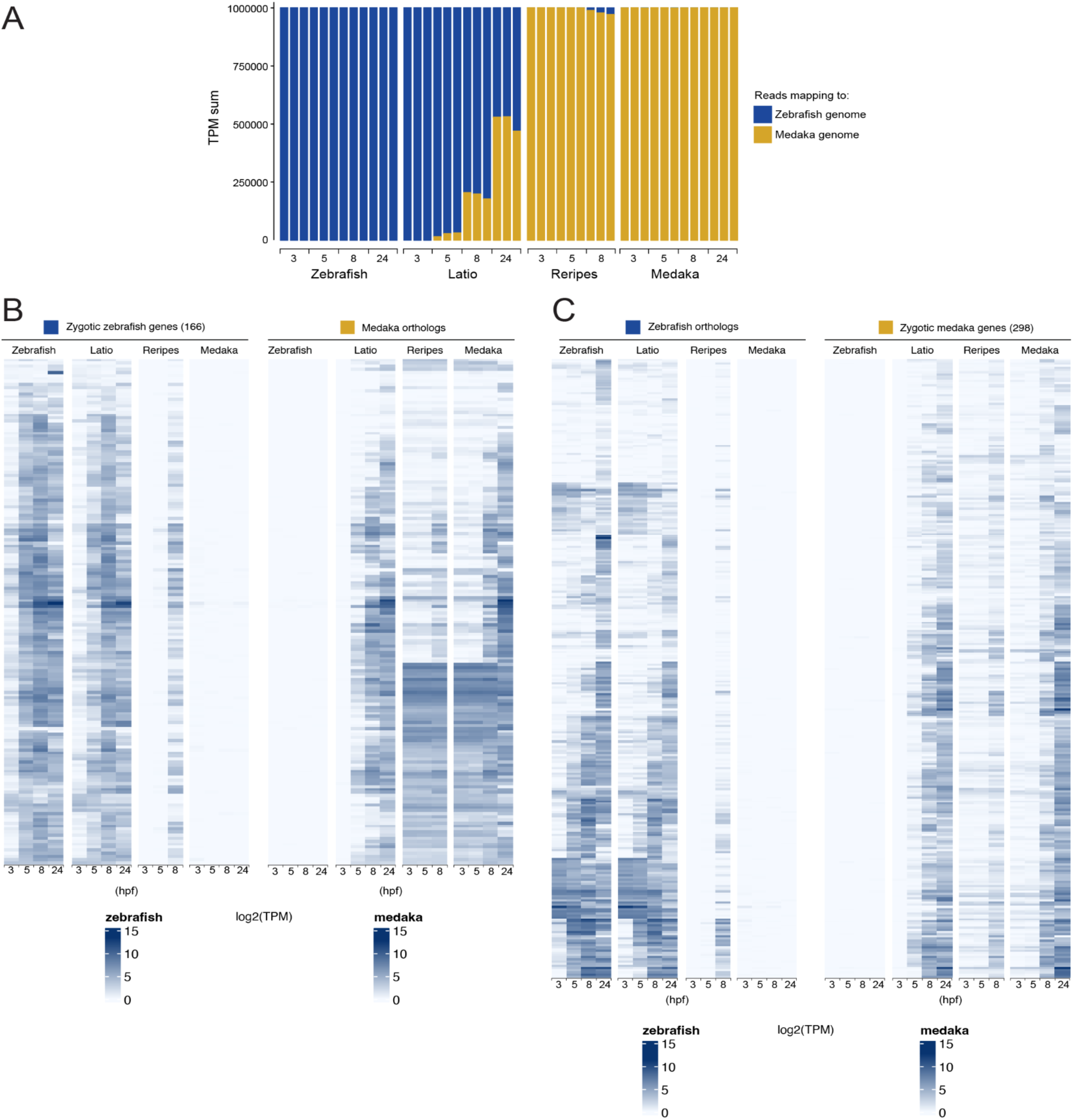
(accompanying Fig 3) **(A)** Total TPM (transcripts per million) mapped to the zebrafish (blue) and medaka (dark yellow) genomes. The first medaka-derived transcripts are detected at 5 hpf in the latio hybrid, whereas zebrafish transcripts are not detected until 8 hpf in reripes. **(B-C)** One-to-one orthologs of zygotic genes in zebrafish and medaka follow the expression dynamics of the maternal ortholog. **(B)** Heat map of gene expression dynamics of the 166 one-to-one orthologs of zebrafish zygotic genes. Log2(TPM) expression values are shown for zebrafish genes (left) and the orthologous medaka genes (right). **(C)** Heat map of gene expression dynamics of the 298 one-to-one orthologs of medaka zygotic genes. Log2(TPM) expression values are shown for zebrafish genes (left) and the orthologous medaka genes (right).

**Supplementary Figure 2.**
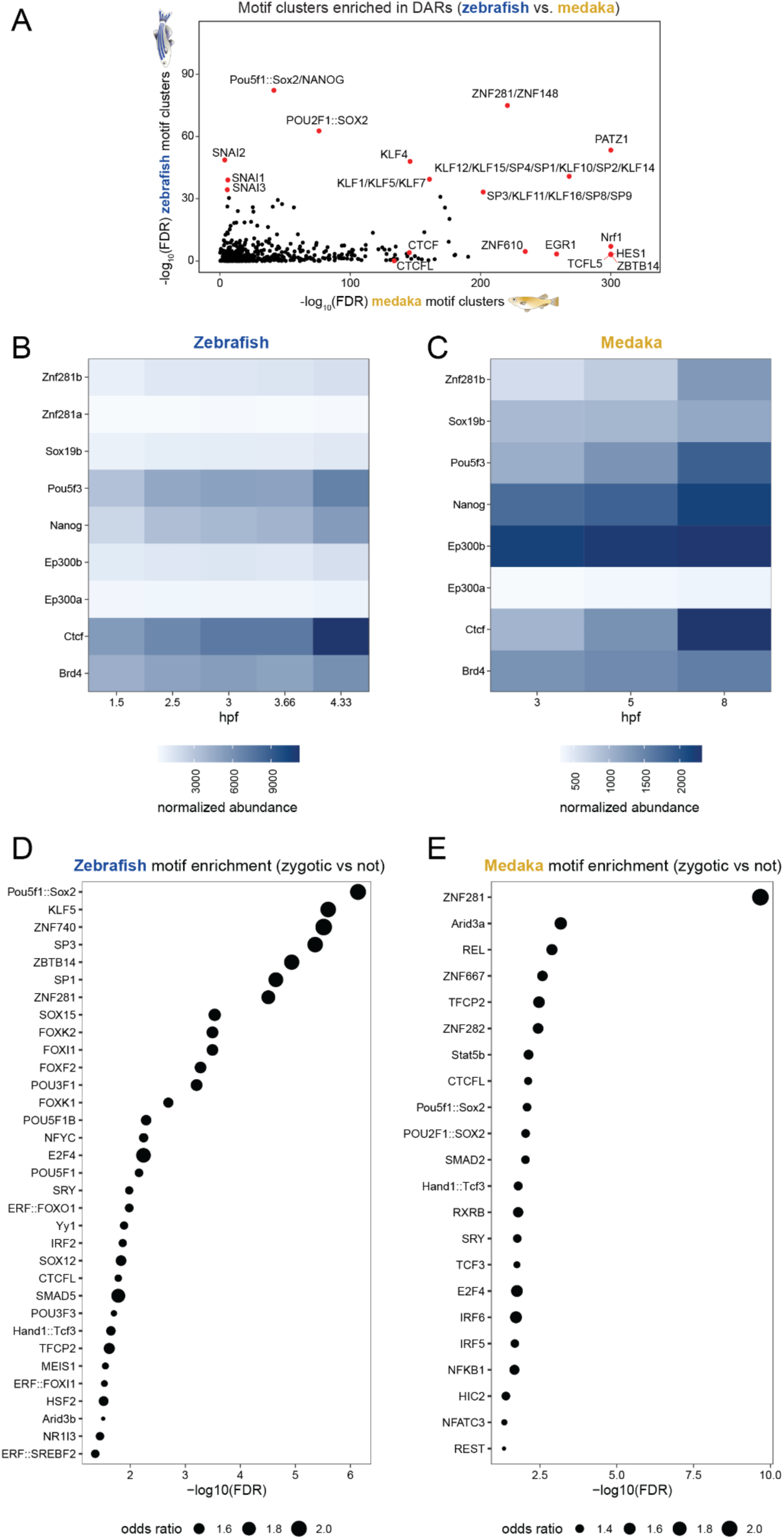
Topmost enriched motifs in differentially accessible zebrafish and medaka genes during ZGA and protein expression fold changes for candidate transcription factors and chromatin remodelers. (accompanying Fig. 5) **(A)** Peak regions significantly increasing in accessibility were used for motif enrichment analysis (see Methods). Motif clusters enriched in zebrafish DARs vs. medaka DARs are plotted. The top 10 motifs in zebrafish and/or medaka DARs are highlighted. **(B-C)** Normalized protein abundance of zebrafish (B) and medaka (C) factors previously implicated in ZGA regulation or corresponding to motifs newly identified as enriched DARs during ZGA. **(D-E)** Most significantly enriched motifs in DARs of zygotic zebrafish (D) and medaka (E) genes compared to non-zygotic genes after filtering for those whose corresponding factors were detected in TMT-MS. Znf281b has the most highly enriched motif in zygotic genes in medaka.

### Supplementary Tables

**Table S1** Expression values from RNA-seq time course in TPM of zebrafish (Z), medaka (M), latio (L) and reripes (R) hybrids at 3, 5, 8, and 24 hpf (no 24 hpf time-point was collected in reripes). Each time-point was collected in triplicates (R1, R2, R3). RNA-Seq data was mapped to a combined zebrafish and medaka genome. Gene set classifications (zygotic, maternal-zygotic, paternal zygotic, unexpressed) are provided in column AV (‘set’).

**Table S2** Orthologous gene list with expression data of zebrafish zygotic genes and their medaka orthologs.

**Table S3** Orthologous gene list with expression data of medaka zygotic genes and their zebrafish orthologs.

## Materials and Methods

### Zebrafish and medaka husbandry

Wild-type medaka fish (*Oryzias latipes,* CAB strain) were raised according to standard protocols (28°C water temperature; 14/10-hour light/dark cycle) and served as wild-type medaka. *Oryzias curvinotus* were obtained from the National Institute for Basic Biology (NIBB) (Okazaki, Japan) and raised under the same conditions as *O. latipes*. Transgenic zebrafish Bouncer-expressing medaka lines were generated as part of this study and are described in detail below. Zebrafish (*Danio rerio*) were raised according to standard protocols (28°C water temperature; 14/10-hour light/dark cycle). TLAB fish, generated by crossing zebrafish AB and the natural variant TL (Tupfel Longfin) stocks, served as wild-type zebrafish for all experiments. *Bouncer* mutant zebrafish and medaka Bouncer-expressing transgenic zebrafish have been published previously^1^. All animal experiments were conducted according to Austrian and European guidelines for animal research and approved by the Amt der Wiener Landesregierung, Magistratsabteilung 58 - Wasserrecht (animal protocols GZ 342445/2016/12 and MA 58-221180-2021-16 for work with zebrafish; animal protocol GZ: 198603/2018/14 for work with medaka).

### Generation of transgenic medaka fish

The ubiquitin promoter and zebrafish *bouncer* N-terminally tagged with sfGFP^1^ were introduced into the pBluescript II SK(-) vector containing I-SceI sites (gift from Katharina Lust) via Gibson cloning. An injection mix containing ubi:sfGFP-zebrafish Bouncer plasmid (10 ng/µL), I-SceI meganuclease (1:10), CutSmart buffer (0.5X), Yamamoto’s ringer’s solution (1X: 1.00 g NaCl, 0.03 g KCl, 0.04 g CaCl_2_·2H_2_O, 0.10 g MgCl_2_·6H_2_O, 0.20 g NaHCO_3_ in 1000 mL, pH 7.3) (0.5X), and phenol red (1:5) was prepared and incubated at room temperature for 1 hour before being placed on ice. Wild-type one-cell medaka embryos were collected from natural medaka crosses and microinjected with 1 nL of injection mix on ice in a medaka injection mold made from 2% agarose in 1X Yamamoto’s ringer solution. Embryos were kept at 28°C and screened for fluorescence 3 or more days post-injection. Fluorescent embryos were reared to adulthood and crossed to wild-type fish to identify founders and generate F1 transgenic fish.

### *In vitro* fertilization with medaka and zebrafish

Wild-type TLAB zebrafish males were set up the night before experimentation with wild-type zebrafish females in a small, plastic breeding tank with a divider separating the two fish. Medaka crosses were set up the night before experimentation inside their tanks in the fish water system with a vertical divider separating one male from two to three females or two males from four to five females. For generation of unfertilized, unactivated medaka eggs, sterile hybrid *O. curvinotus* x *O. latipes* males, for example derived from *O. curvinotus* x *O. latipes* hybrid males^10^, were used to mate with wild-type CAB females or transgenic CAB females expressing zebrafish Bouncer (tg[ubi:sfGFP-zebrafish Bouncer]). On the day of experimentation, sperm was collected from the zebrafish males after anesthetization in 0.1% (w/v) tricaine (25X stock solution in dH_2_O, buffered to pH 7-7.5 with 1 M Tris pH 9.0) in fish system water. A capillary fitted with small plastic tubing and a pipette filter tip on the other end was used to mouth-pipette sperm from the urogenital opening of each male positioned belly-up in a slit in a sponge wetted with fish water. Sperm was transferred directly to a 1.5-mL tube containing Hank’s balanced salt solution (see below) on ice. In general, based on the number of clutches to be fertilized, one male was used per 100 µL of Hank’s saline. Because sperm is used in great excess during IVF, any concentration above 50,000 sperm/ul was used.

Freshly spawned medaka eggs were collected directly from the bodies of females using a net with fine mesh and by gently pulling the eggs from the fish in the net using the thumb and forefinger on the outside of the net, taking care not to crush any eggs during removal. Collected eggs were placed into petri dishes containing 1X Yamamoto’s ringer’s solution. After collection, eggs were visually inspected under a dissection microscope to remove any crushed or already activated eggs. As much ringer’s solution as possible was removed from each dish such that the eggs remained submerged when the dish was tilted on its lid. 45 µL of sperm suspension was pipetted directly onto the eggs. After 2 minutes, 2-3 µL of 0.1% (w/v) calcimycin in DMSO was pipetted carefully onto the eggs. After 10 minutes, the dishes were filled with E3 medium (5 mM NaCl, 0.17 mM KCl, 0.33 mM CaCl_2_, 0.33 mM MgSO_4_, 0.00001% methylene blue) or 1X Yamamoto’s ringer’s solution before being placed into an incubator at 28°C.

### Confocal imaging of zebrafish and medaka sperm in the medaka micropyle

Zebrafish sperm and medaka eggs were collected as described above for IVF. To label sperm, MitoTracker Deep Red (1:400) was added to Hank’s solution before sperm collection. Unactivated medaka eggs (one per fertilization movie) were placed into a medaka injection mold made with 2% agarose in 1X Yamamoto’s ringer solution in a standard petri dish and positioned with a metal probe under a dissection microscope so that the micropyle was visible. The mold containing the positioned egg was then placed onto an LSM800 Examiner Z1 (Zeiss) upright confocal microscope and the egg was imaged with a 20x/1.0 plan-apochromat water objective. Imaging was started and 6 million sperm were pipetted as close as possible to the egg under the objective. For experiments with calcimycin, 2-3 µL of 0.1% calcimycin in DMSO was pipetted as near to the egg as possible 1.5-2 minutes (such that sperm had reached the end of the micropyle) post sperm addition. During imaging, the micropyle was in focus at all times until sperm fusion and sperm were followed by manually adjusting focus as they moved through the micropyle and if they fused with the egg.

### Bright field imaging

Live embryos were imaged in their chorions in 1.5% methylcellulose on a glass slide using a dissection microscope (Stemi 508, Zeiss) at 4X magnification using FlyCapture2 (Point Grey Research) and a BlackFly USB color camera (BFLY-U3-23S6C-C).

### rRNA-depleted RNA library preparation and sequencing

Embryo samples for latio hybrids were generated as previously described^1^. Wild-type medaka CAB and zebrafish TLAB embryos were collected from natural crosses after being separated the night before sample collection. Reripes samples were generated as described above. All samples were collected in triplicates at each time point (3, 5, 8, and 24 hpf (except 24 hpf for reripes hybrids). To isolate RNA, samples of 10 embryos per time point were homogenized in TRIzol with an electric pestle and flash-frozen in liquid nitrogen. RNA isolation was performed using standard protocols (phenol/chloroform extraction followed by isopropanol precipitation). RNA concentration was measured using a Fragment Analyzer System (Agilent). rRNA depletion was performed using either the RiboCop rRNA Depletion Kit (Lexogen) using 500 ng of RNA per sample (zebrafish, latio and medaka samples) or the Ribo-zero Gold Kit (Illumina, reripes samples). Libraries were prepared using NEBNext Ultra Directional RNA Library Prep Kit for Illumina (NEB) (zebrafish, latio and medaka samples) or the Illumina TruSeq Stranded Total RNA Library Prep kit (reripes samples) and checked using a Fragment Analyzer System (Agilent) before multiplexing. Sequencing was performed on Illumina HiSeqV4 SR100 and NovaSeq 150 bp paired-end sequencing platform.

### RNA-seq analysis

RNA-seq raw reads were adapter trimmed using bbmap v38.26 and mapped to a *D.rerio* and *O. latipes* hybrid reference genome composed of the Ensembl build 96 genomes GRCz11 and OlASM223467 using Hisat2 v2.1.0. Reads in genes were counted using htseq v0.11.0 (-m intersection-nonempty). TPM estimates were derived from gene-level counts by normalizing for gene length, sequencing depth, and scaling to the sum of 1 million. Only uniquely mapping reads were used for analysis (genes for which reads mapped to the foreign genome in the purebred fish species with an expression level TPM > 5 were filtered out). Orthologous genes were obtained from Ensembl Compara with homology type ortholog_one2one. *D. rerio* zygotic gene candidates were defined based on a SLAMSeq embryonic time course^22^. Candidate *O. latipes* zygotic genes were defined as genes that are not detected at stage 6 (TPM<0.001) but are detected at stage 11 (TPM>2) based on GSE136018^25^.

### Tandem mass tag mass spectrometry (TMT-MS) sample preparation

Wild-type medaka embryos were obtained via IVF using wild-type medaka sperm and eggs from females mated with infertile hybrids, as described above, to ensure embryo synchronization. Embryos were incubated at 28°C until the desired time of collection (3, 5, or 8 hpf). To dissect animal caps for proteomic analysis, embryos were rolled on fine sandpaper and pipetted into a Silguard dish filled with 1X Danieau’s solution (58 mM NaCl, 0.7 mM KCl, 0.4 mM MgSO_4_•7H_2_O, 0.6 mM Ca(NO_3_)_2_, 5 mM HEPES) at 28°C. Embryos were cut in half laterally or an opening was cut in the chorion using fine microdissection scissors; animal caps were manually dissected from the yolk and transferred to a second dish of 1X Danieau’s and collected in a tube on dry ice in a small volume of 1X Danieau’s. Samples for each time point were collected in 4 replicates.

### Filter-aided sample preparation (FASP) of medaka embryo caps

Samples were processed using the FASP method with minor modifications^54^. Briefly, embryo caps were dissolved in 30 µl of SDT buffer (4% SDS, 0.1M DTT in 0.1M TEAB; pH 7.5) and incubated for 10 min at 95°C. The samples were sonicated with an ultrasonication probe using the Ultrasonic processor UP100H (Hielscher) for 30 cycles (0.5 s, 50% amplitude). Protein concentration was determined by measuring the fluorescence of tryptophan. Protein extracts were mixed with 200 µl of UA solution (8 M urea in 0.1 M TEAB, pH 8.5), transferred to Microcons (MWCO 30 kDa), and centrifuged (20 min, 14,000 × g, RT). After washing with 200 µl of UA solution, proteins were alkylated by adding 100 µl of 0.1 M iodoacetamide in UA solution and incubated for 20 min at RT in the dark. This was followed by centrifugation at 14,000 × g for 15 min at RT. Thereafter, the ultrafiltration unit was washed twice with 100 µl of UA solution and twice with 100 µl of 50 mM TEAB. Proteins were digested directly in the ultrafiltration unit by adding 50 µl of TEAB buffer containing trypsin (Mass spectrometry grade; Promega) in a 1:20 ratio (protease: protein, w/w). After overnight digestion at 37°C, peptides were eluted into fresh collection tubes by centrifugation at 14,000 × g for 15 min at RT. The ultrafiltration unit was then washed once by addition of 50 µl 50-mM TEAB buffer and centrifuged for 10 min at 14,000 × g. To evaluate the digestion efficiency and quantify the peptide yield, 100 ng of each sample was analyzed on a monolithic HPLC system.

### Labelling with Tandem Mass Tags (TMT)

Approximately 50 µg (in 100µl of 100 mM HEPES, pH 7.6) of each sample was labelled with one separate channel of the TMTpro 16plex Label Reagent (ThermoFisher) according to the manufacturer’s description. The labelling efficiency was determined by LC-MS/MS with a small aliquot of each sample. Samples were mixed in equimolar amounts and equimolarity was again evaluated by LC-MS/MS. The mixed sample was acidified to a pH below 2 with 10% trifluoroacetic acid (TFA) and was desalted using Sep-Pak C18 1 cc Vac cartridges (50 mg, Waters). Peptides were eluted with 3 x 150 µl 80% acetonitrile (ACN) in 0.1% formic acid (FA), followed by freeze-drying.

### Strong cation exchange (SCX) separation and mass spectrometry analysis

The dried sample was dissolved in 70 µl of SCX Buffer A (5 mM NaH_2_PO_4_, pH 2.7, 15% ACN) and 200 µg of peptide were loaded on the column. SCX was performed using a custom-made TSKgel SP-2PW SCX column (5 µm particles, 12.5 nm pore size, 1 mm i.d. x 250 mm, TOSOH Bioscience) on an UltiMate System (ThermoFisher Scientific) at a flow rate of 35 µl/min. For the separation, a ternary gradient was used. Starting with 100% buffer A for 10 min, followed by a linear increase to 10% buffer B (5 mM NaH_2_PO_4_, pH 2.7, 1 M NaCl, 15% ACN) and 50% buffer C (5 mM Na_2_HPO_4_, pH 6, 15% ACN) in 10 min, to 25% buffer B and 50% buffer C in 10 min, to 50% buffer B and 50% buffer C in 5 min and an isocratic elution for an additional 15 min. The flow-through was collected as a single fraction, and the gradient fractions were collected every minute. 60 fractions were collected; the first 20 fractions were discarded and the low abundance fractions were pooled. ACN was removed by vacuum centrifugation and the samples were acidified with 0.1% TFA and analyzed by LC-MS/MS.

The nano HPLC system used was an UltiMate 3000 HPLC RSLC nano system (Thermo Scientific) coupled to an Exploris 480 mass spectrometer, equipped with a FAIMS pro interface and a Nanospray Flex ion source (all parts ThermoFisher Scientific). Peptides were loaded onto a trap column (ThermoFisher Scientific, PepMap C18, 5 mm × 300 μm ID, 5 μm particles, 100 Å pore size) at a flow rate of 25 μL/min using 0.1% TFA as the mobile phase. After 10 min, the trap column was switched in line with the analytical column (ThermoFisher Scientific, PepMap C18, 500 mm × 75 μm ID, 2 μm, 100 Å) operated at 30°C. Peptides were eluted using a flow rate of 230 nL/min, starting with the mobile phases 98% A (0.1% FA in water) and 2% B (80% ACN, 0.1% FA) and linearly increasing to 35% B over the next 120 minutes, followed by an increase to 90% B in 5 minutes, where it stayed for 5 minutes before changing back to 98% A for equilibration. The Exploris 480 mass spectrometer was operated in data-dependent mode, performing a full scan (m/z range 350-1200, resolution 60,000, target value 1E6) at 3 different compensation voltages (CV -40, -55, -70), each followed by MS/MS scans of the most abundant ions for a cycle time of 1 second per CV. MS/MS spectra were acquired using an HCD collision energy of 34%, isolation width of 0.7 m/z, resolution of 45.000, first fixed mass 110, target value at 2E5, minimum intensity of 2.5E4 and maximum injection time of 120 ms. Precursor ions selected for fragmentation (including charge state 2-6) were excluded for 45 s. The monoisotopic precursor selection (MIPS) mode was set to “peptide”, and the “exclude isotopes” and “single charge state per precursor” features were enabled.

### Proteomics data analysis

For peptide identification, the RAW files were loaded into Proteome Discoverer (PD, version 3.0.0.757, ThermoFisher Scientific). All MS/MS spectra were searched using MS Amanda v2.0.0.19742^55^. Trypsin was specified as a proteolytic enzyme cleaving after lysine and arginine (K and R) without proline restriction, allowing for up to 2 missed cleavages. Mass tolerances were set to ±10 ppm at the precursor and ±10 ppm at the fragment mass level. The RAW files were searched against the ENSEMBL database, using taxonomy Medaka/Oryzias_latipes (release 2022_12; 36,075 sequences; 21,697,622 residues), with common contaminants appended. Iodoacetamide derivative on cysteine was set as a fixed modification; deamidation on asparagine and glutamine, oxidation on methionine, carbamylation on lysine, TMTpro-16plex tandem mass tag on lysine as well as carbamylation on peptide N-Terminus, TMTpro-16plex tandem mass tag on peptide N-Terminus, glutamine to pyro-glutamate conversion at peptide N-Terminus and acetylation on protein N-Terminus were set as variable modifications. Results were filtered for a minimum peptide length of 7 amino acids and 1% FDR at the peptide spectrum match (PSM) and the protein level using the Percolator algorithm^56^ integrated in Proteome Discoverer. Additionally, an Amanda score of at least 150 was required. The localization of the modification sites within the peptides was performed with the tool ptmRS, based on the tool phosphoRS^57^. Identifications were filtered using the filtering criteria described above, including an additional minimum PSM-count of 2 for the summed-up fractions. Peptides were quantified based on Reporter Ion intensities extracted by the “Reporter Ions Quantifier”-node implemented in Proteome Discoverer. Proteins were quantified by summing unique and razor. Proteins were filtered to be identified by a minimum of 3 quantified peptides. Protein abundance normalization was done using sum normalization. Statistical significance of differentially expressed proteins was determined using limma^58^.

### ATAC-sequencing sample and library preparation

Both purebred (medaka and zebrafish) and hybrid (latio and reripes) embryos were generated using IVF as described above. Zebrafish and latio hybrid embryos were dechorionated with pronase (1 mg/ml) treatment upon reaching the desired time post-fertilization and manually de-yolked using fine forceps in 1X PBS at 28°C. To reach a sufficient number of cells for each time point, 60, 15, and 10 embryo caps were collected at 3, 5, and 8 hpf, respectively, on ice. Medaka and reripes hybrid embryos were mechanically dechorionated in Leibovitz’s L-15 medium (Gibco) inside a small plastic bag by crushing with a metal cooling block. The medium was then collected and centrifuged at 300 x g for 10 minutes at 4°C. The bag was washed twice with 1 mL of L-15 medium and this was added to the first volume of cells, which was centrifuged each time and the supernatant removed. Embryonic cells were resuspended in lysis buffer (10 mM Tris HCl, 10 mM NaCl, 3 mM MgCl2, 0.1% NP-40, 0.1% Tween20, and 0.01% digitonin, pH 7.5) and incubated on ice for 5 minutes. Dilution buffer (10 mM Tris HCl, 10 mM NaCl, 3 mM MgCl2, 0.1% NP-40, 0.1% Tween20, pH 7.5) was added and the cells were resuspended before centrifugation at 500 rcf for 10 minutes at 4°C. After removal of the supernatant, the lysis and dilution steps were repeated and the supernatant was removed again. Tn5 transposase (Illumina) and TD buffer (Illumina) were added directly to the lysed nuclei and incubated for 30 minutes at 37°C with shaking (500 RPM). The transposition reaction was cleaned up immediately using the Zymo DNA Clean and Concentrator Kit. Library amplification was done using the purified, tagmented DNA, forward and reverse primers (25 uM) provided from the NGS facility at the Vienna BioCenter, EvaGreen® dye (Biotium) and Q5® High-Fidelity 2X Master Mix (NEB, M0492S).

### ATAC-seq analysis

ATAC-seq reads were quality trimmed using trimmomatic v0.39 using parameters ILLUMINACLIP:NexteraPE-PE.fa:2:30:10:2:TRUE HEADCROP:10 LEADING:3 TRAILING:3 MINLEN:36. Trimmed reads were aligned to a hybrid genome of *O. latipes* (ASM223467v1) and *D. rerio* (GRCz11) Ensembl version 96 using bowtie v2.4.4 with parameters --dovetail --no-mixed --no-discordant. Alignments were deduplicated and quality filtered using samtools v1.15.1 at MAPQ>=30 and discarding any secondary alignments. Reads mapping to mitochondrial genomes were removed prior to normalization and further analyses. Normalized bigwig files were generated using deeptools v3.5.1 with parameters -bs 20 –normalizeUsing RPKM. For peak calling, ATAC-seq read pairs with insert sizes > 150bp were discarded. Peaks were called using macs2 with parameters --format BAMPE -q 0.05 --call-summits, pooling replicates. Overlapping peaks were merged and compiled into a consensus matrix for comparison across all samples. This matrix was further filtered to only include consensus peaks for which at least one sample had a q-value smaller or equal to 1e-7. Promoter regions were defined as a window 2-kb upstream and 0.5-kb downstream of all annotated TSSs. For the *D. rerio* genome standard Ensembl v96 annotations were used. To improve annotations for the *O. latipes* genome, alternative TSS-annotations according to GSE136018 SI were used and mapped to Ensembl gene IDs (mapping table available at: http://tulab.genetics.ac.cn/medaka_omics/). In case of ambiguous mappings (more than one alternative annotation) we kept the one with the shortest distance to the original TSS. Reads overlapping promoters/peaks were counted using featureCounts (Rsubread package). Counts based on reads aligned to the respective other genome in purebred samples were set to zero. In cases where a gene showed an average of 10 reads misaligned across purebred samples, the corresponding gene/region was blacklisted. TSS/promoter enrichment of reads over the genome was calculated as the fraction of the total number of reads overlapping promoter regions normalized to total promoter length and the total number of reads mapped normalized to genome length. FPKM normalization for heatmaps and metaplots was performed separately for each genome in each sample. Heatmaps and metaplots were calculated and plotted in R using standard plotting libraries (ggplot).

### Transcription factor motif analysis

A consensus set of ATAC-seq peak regions was compiled by merging overlapping peak regions across samples. ATAC-seq fragments were counted over these consensus peaks and differentially accessible regions (DARs) were called separately for each genome using the R-package DESeq2 with foldchange shrinkage. P-values were adjusted for multiple testing using the Benjamini-Hochberg method. Peak regions that were significantly increasing in accessibility (log2 fold change >1, FDR <5%) between 5 and 8 hpf for medaka and between 3 and 5 hpf for zebrafish, respectively, were selected for motif enrichment analysis. As background sequences, an equally large and non-overlapping set of peaks with the lowest fold changes (closest to 0) were selected. Transcription factor matrices in the JASPAR 2022 core vertebrate database were supplemented by a set of nanog motif matrices from HOCOMOCOv10 using the R-package MotifDb. This set of motif matrices was then scanned against the sequences underlying the selected peak regions using the motifmatchR package, reporting any hits at a significance threshold of 0.0001. Significance of motif occurrence in the differentially accessible regions (DARs) was evaluated against the background sets using fisher tests, correcting for multiple testing (Benjamini-Hochberg). Results were compared between zebrafish and medaka DARs, yielding a number of transcription factors that were either preferentially enriched in zebrafish, medaka, or both. For alternative analyses (SI), motif matrices were additionally clustered and sufficiently similar motifs (pearson correlation >= 0.95) were merged using the universalmotif R-package to reduce motif redundancy.

